# Perceptual awareness of near-threshold tones scales gradually with auditory cortex activity and pupil dilation

**DOI:** 10.1101/2024.03.14.584980

**Authors:** Laura Doll, Andrew R. Dykstra, Alexander Gutschalk

## Abstract

Perceptual awareness covaries with negative-going responses in sensory cortex, but the derived concept of perceptual awareness negativity has been criticized a.o. because of its presence for undetected stimuli. To evaluate this objection, we combined magnetoencephalography, electroencephalography, and pupillometry to study the roles of sustained attention and response criterion on the auditory awareness negativity. Participants first detected distractor sounds and denied hearing task-irrelevant near-threshold tones, which evoked neither awareness negativity nor pupil dilation. These same tones evoked responses when task-relevant, stronger for hit but also present for miss trials. To explore if response criterion could explain the presence of responses for miss trials, participants rated their perception on a six-point scale. Decreasing perception ratings were associated with gradually reduced evoked responses, consistent with signal detection theory. These results support the concept of an awareness negativity that is modulated by attention, but does not exhibit a non-linear threshold mechanism.

## Introduction

Sensory cortex plays a central role in perceptual awareness. Several studies have demonstrated that a negative-going response recorded in MEG and EEG – referred to as perceptual awareness negativity – closely corresponds to behaviorally confirmed awareness across sensory modalities^1,2^. However, there is ongoing debate about whether sensory cortex itself subserves perceptual awareness^3,4^ or if sensory cortex is instead the final processing stage to transmit information to a higher-order network including fronto-parietal cortex that enables perceptual awareness^5,6^. Furthermore, the neural machinery for perceptual awareness also includes subcortical networks: Ascending monaminergic projections from dorsal pons and midbrain are required to maintain a state of wakefulness and sustained attention^7–9^, attentional mechanisms of the thalamus may be important^10,11^, and sensory detection in a mouse model has been shown to involve projections from sensory cortex to subcortical targets^12^.

Perceptual awareness is often studied with stimuli that are near-threshold or masked. Within the framework of signal detection theory (SDT)^13^, the detection of a stimulus depends first on the strength of a hypothetical, normally distributed neural representation of the stimulus, and second on the decision criterion set by the participant. Despite the long history of SDT in perception research and how central the question is to the neural basis of perceptual awareness, it has not been firmly established on which neural signal the perceptual decision is based. Potential sources of the variability associated with perceptual awareness of a near-threshold stimulus are (i) the processing along the ascending sensory pathway, e.g. in sensory cortex^14^, (ii) the propagation of the signal from sensory to frontal cortex^15^, or (iii) networks related to salience and attention^16,17^. In the present study, we explored if the AAN could embody the variable neural representation predicted by SDT In parallel to M/EEG, we recorded the pupil dilation response (PDR) as an additional index of networks underlying arousal and sustained attention^18^.

First, we probed how the AAN is modulated by attention and task relevance; listeners had to indicate their detection of near-threshold tones via a button press or perform a competing auditory task to draw attention away from the same near-threshold tones. Next, participants rated their perception of near-threshold tones on a six-point scale, which allowed us to investigate the relationship between neural response and strength of perception, both in the range considered detected and missed in the binary response scheme of the first experiment.

## Results

### Task-related modulation of near-threshold tones

To explore the attention- and task-dependent modulation of perceptual awareness, three identical sequences of near-threshold (NT) pure tones were presented in continuous white noise, together with more salient, transient amplitude-modulation (AM) epochs of the noise. In the first run, participants had to detect the AM noise and were not informed about the NT tones. In the second run, the task was to detect the NT tones and ignore the AM-noise epochs (Fig. 1A). In the third run, participants were again instructed to detect the AM noise, similar as in run 1. We expected that the participants were not aware of the NT tones in the first run, but that they hear more tones in the third run, after actively detecting them in the second run, similar to previous experiments^19,20^.

**Fig. 1.**
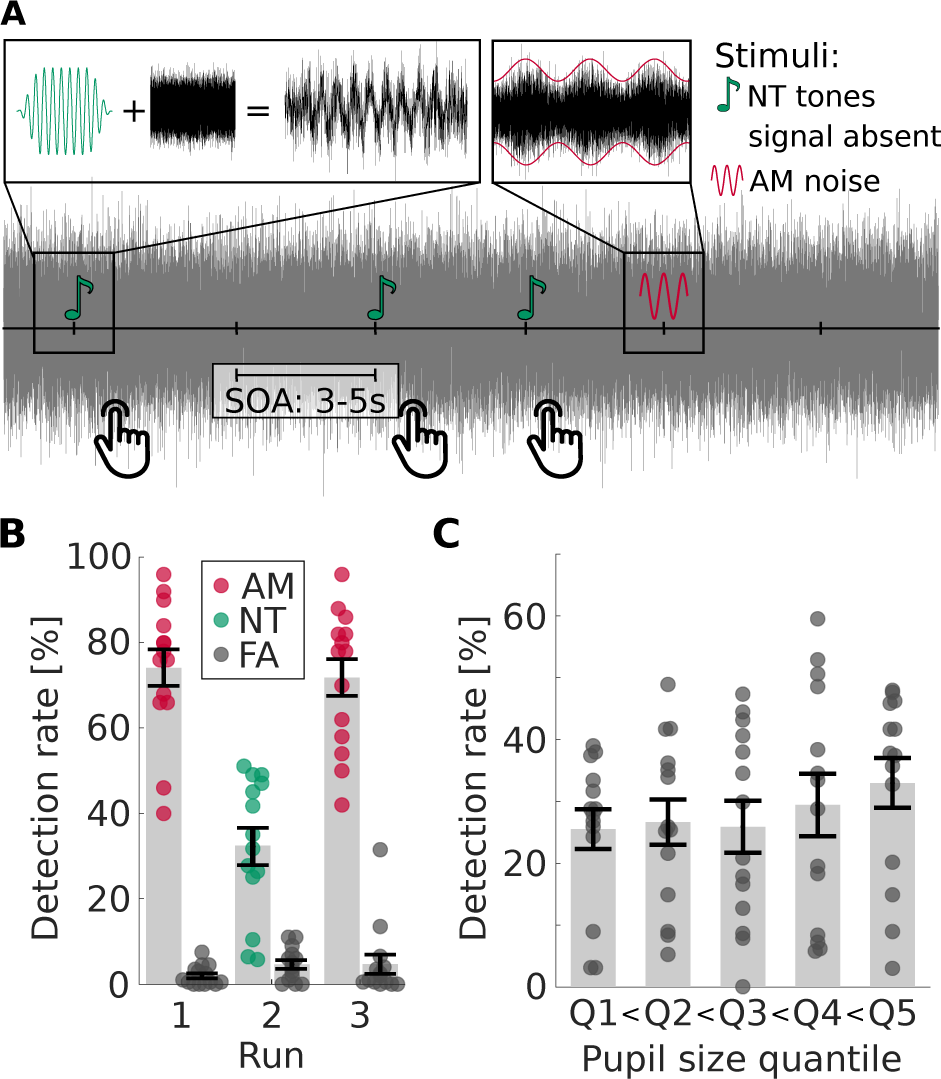
Paradigm and behavioral results of Experiment 1 (Selective attention task). A) Experimental paradigm: transient AM noise, NT tones, and subthreshold tones (i.e., signal-absent trials) are presented in continuous noise. A button press indicates detected targets, i.e. AM noise in runs 1 and 3, NT tones in run 2. B-C) Behavioral results. Single points represent the respective value of a single participant, bar height and error bars represent the group mean and standard error. B) Detection and false alarm rate for the three runs. C) Detection rate of NT tones in the second run as a function of baseline pupil size.

### Behavior

On average, more than 70% of the AM noise was detected in runs 1 and 3, respectively, while the detection rate for NT tones in run 2 was only around 35% (Fig. 1B). False-alarm rates were low in all runs. Decision criteria were 0.8 ± 0.3 and 0.7 ± 0.4 (mean ± s.d.) in runs 1 and 3 for detection of AM noise and 1.2 ± 0.4 in run 2 for detection of NT tones, showing that participants were more conservative during the detection of NT tones. This was confirmed by a paired t-test (mean criterion of run 1 and 3 vs. criterion of run 2; t_13_=-4.8, p=0.0004). When asked regarding their perception of NT tones after run 1, no participant reported having heard them. When asked again after run 3, 7 out of 14 participants reported having heard at least some NT tones in the last run of the experiment. While participants were not systematically asked about hearing AM noise in run 2, a number of participants spontaneously reported being distracted by the interfering AM noise.

Overall, detection rates of the NT tones were somewhat lower than expected (50% for NT tones) based on pilot measurements with the adaptive testing procedure (see Method section and ^21^). Several factors might contribute to this difference: First, adaptive tests are usually performed several times to provide a stable estimate of the detection rate. This was not done here in order to keep the total duration of the experiment within reasonable limits. Second, the main experiment requires sustained attention over a longer period of time, while the adaptive test takes only a few minutes. Supporting the role of sustained attention, splitting run 2 into five parts showed a decreasing detection rate as a function of time (mean ± s.d. = 35 ± 18, 39 ± 21, 30 ± 18, 33 ± 20, 21 ± 17 % for each part, respectively), confirmed by repeated-measures analysis of variance (rmANOVA) (F_4,52_=4.16, p=0.016). In addition to the longer duration of the experiment, the time interval between targets was also increased, leading to a higher demand on vigilance and sustained attention. Third, the presence of AM might impair the NT detection, as it was reported by some participants to be distracting.

Evaluation of the detection rate as a function of pre-stimulus pupil size revealed a small increase of detection rate with pupil size (Fig. 1C). A linear contrast analysis showed a significant linear (F_1,13_=12.62, p=0.0035) but no quadratic effect (F_1,13_=1.09, p=0.32). These findings are in contrast to previous publications^8,22^ reporting an inverted U-shaped relationship (i.e., a quadratic effect) between arousal and performance measured either by reaction time^22^ or accuracy^8^. Our participants may not have reached high levels of arousal during experiment 1, which would suggest that we see only part of a possibly U-shaped curve.

### M/EEG

As expected, combined M/EEG source-mapping showed negative-going activity in auditory cortex (AC) around 200 ms post-stimulus onset and additional activity in the region of retrosplenial and posterior cingulate cortex (RSC/PCC) around 500 ms post-stimulus onset for detected targets (Fig. 2). For missed NT tones, there was almost no activity at 200 ms, and the activity at 500 ms was mostly restricted to AC.

**Fig. 2.**
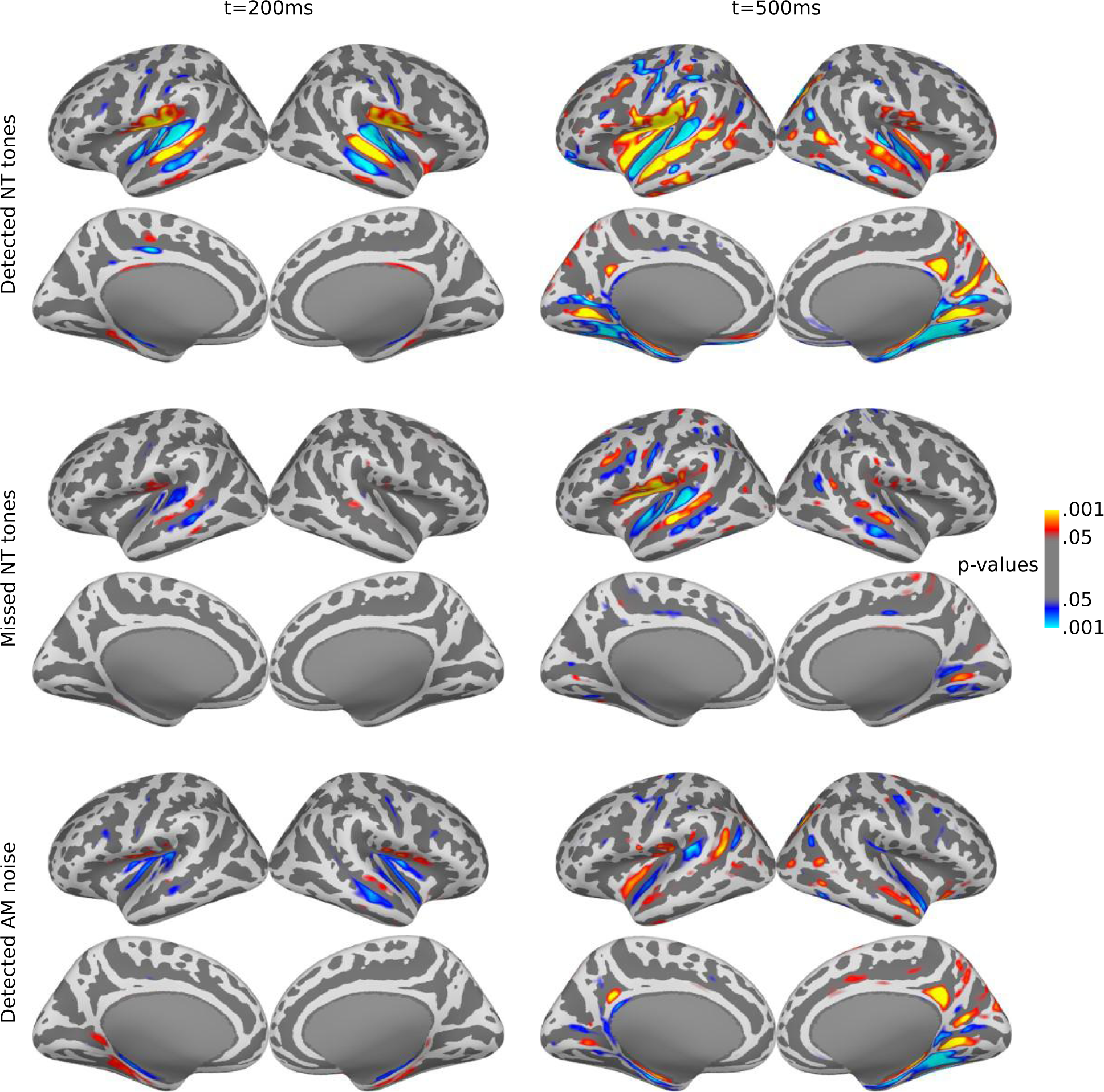
dSPM maps for detected targets of Experiment 1 (Selective attention task). Left column: t=200ms, right column: t=500ms. Two rows each belong to the following conditions: detected NT tones, missed NT tones, and AM noise. The passive stimuli and the undetected AM noise did not show any relevant activity, and are therefore not shown. Since AM-noise maps of runs 1 and 3 showed no visible difference, they are combined here.

For the comparison between conditions, activation time courses were calculated based on anatomical regions of interest (ROIs). Because of the variable hit rates, this analysis was based on the minimum norm estimate (MNE), a linear estimate whose amplitude does not scale with number of trials. In AC (Fig. 3, left column), detected NT tones (run 2) evoked prominent negative-going activity starting around 150 ms and sustaining until 700 ms after stimulus onset before slowly returning to baseline. Missed NT tones in run 2 also elicited a similar but much smaller response. Significance of the responses to both NT hits and NT misses were confirmed with permutation cluster tests^23^ in the time interval 100-400 ms (Table S1). In contrast, no significant negative-going response for NT tones was observed in runs 1 and 3, when they were not task-relevant. When the data were analyzed separately between participants who heard or didn’t hear NT tones during the last run, a small, non-significant negative deflection is observed in auditory cortex, but only for listeners who heard the tones (Fig. S1). One potential factor for the lack of significance could be a small number of tones perceived; one participant retrospectively estimated the number of detected tones to be 50% of those in the previous run; another reported hearing only 3-4 tones. This analysis also reveals that the subgroup of participants who did not perceive the NT tones in run 3 show a positive deflection for NT tones in the last run, similar to both groups in the first run (Fig. S1). This could indicate the presence of small P1 and P2 responses irrespective of perception that are canceled by negative-going components when the stimuli are perceived or attended.^24^

**Fig. 3.**
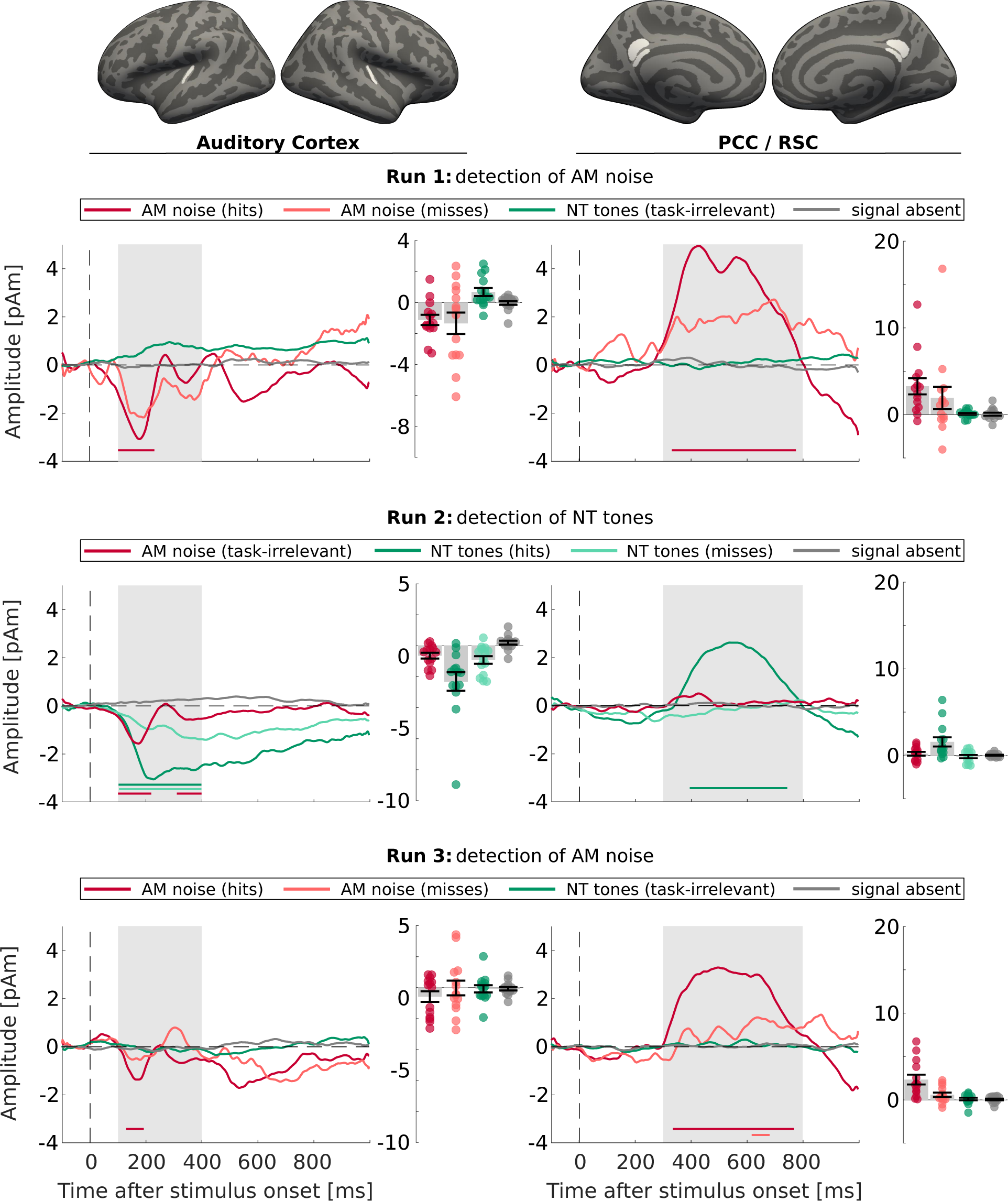
M/EEG waveforms of Experiment 1 (Selective-attention task). Waveforms extracted from the MNE-based source estimate in bilateral AC (left) and RSC/PCC (right) together with mean amplitudes in the time windows of interest (gray background in the waveform plots, AC: 100-400ms, RSC/PCC: 300-800ms). Rows 1-3 show the results of the three runs, respectively. The colored horizontal bars in the waveform plots mark the time windows of significant activity (permutation cluster test). In the amplitude plots, each circle represents the respective amplitude of one participant, bar height and error bars represent the group mean and standard error. H/MAM: hits/misses AM noise, NT: task-irrelevant NT tones, H/MNT: hits/misses NT tones, signal absent: signal-absent trials without button press (correct rejections).

A different pattern was observed for the AM noise. Here, a negative-going transient was observed in AC in all three runs, but was not significant for the missed AM noise trials in runs 1 and 3. Comparison of the response amplitude across all stimuli, irrespective of detection, revealed no significant difference between the three runs (permutation cluster tests, no clusters found for run 1 vs. run 2; non-significant cluster for run 1 vs. run 3 and run 2 vs. run 3: t_1-3_=100-141 ms, p_1-3_=0.08; t_2-3_=100-101 ms, p_2-3_=0.21). The same analysis for NT tones revealed significant differences between all runs (t_1-2_= 100-400 ms, p_1-2_=0.00004; t_2-3_=111-400 ms, p_2-3_= 0.0007; t_1-3_=177-326, p_1-3_=0.027).

The ROI analysis in the RSC/PCC showed a strong and significant response consistent with a P3^25^ from approximately 400 to 600 ms for detected target stimuli in all three runs, and for undetected AM noise in run 3 (Fig. 3, right columns), but not for the other missed targets or for stimuli that were not task-relevant.

### Pupil dilation response

The pupil dilation response (PDR) started at approximately 500 ms, peaked first at 1000 ms and again around 1750 ms, and decreased slowly afterwards (Fig. 4). To focus on the onset of the PDR response, we evaluated its first derivative, or PDR’ (time interval 500 – 1000 ms used for analysis). The pupil showed a strong dilation after all task-relevant detections. We also observed a small PDR for missed target stimuli. Although the maximum amplitude is only slightly larger than the one for signal-absent trials, the first derivative shows that the PDR’ for signal-absent trials is a rather sustained offset, while the response to missed targets results in a notable peak with latencies similar to the PDR’ of detected targets. In contrast, we did not observe a significant PDR for stimuli that were not task-relevant, including AM noise; clusters found for task-irrelevant stimuli in runs 2 and 3 did not reach statistical significance.

**Fig. 4.**
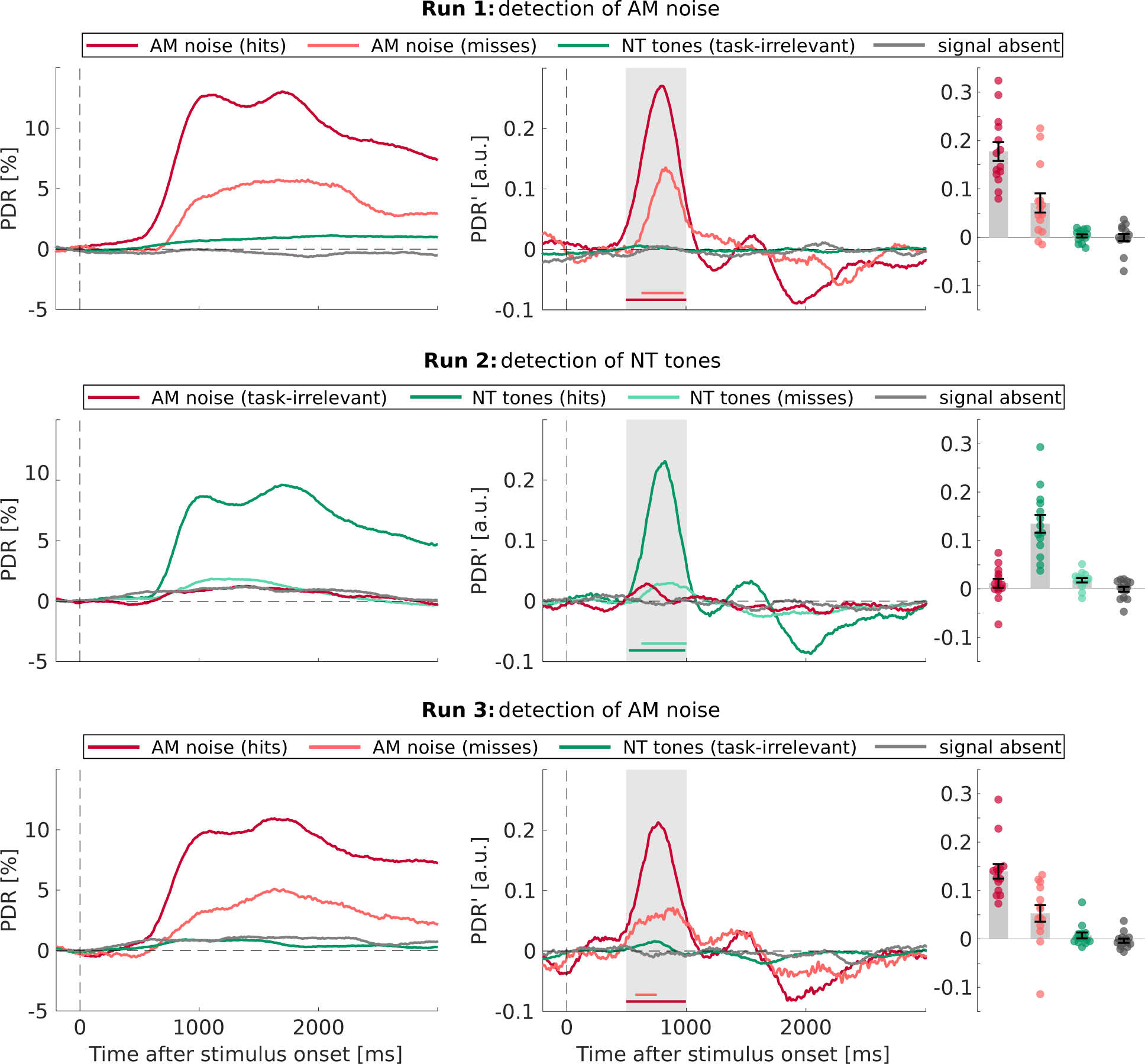
Pupil dilation response of Experiment 1 (Selective attention task). PDR (left column) and its first derivative (PDR’, middle column) for the three runs (rows 1-3, respectively) of Experiment 1. Statistical analysis was conducted on PDR’ from 500 to 1000 ms (gray background); colored horizontal bars mark the time windows of significant activity (permutation cluster test). Amplitudes averaged across the 500 ms time window are plotted on the right. Each circle represents one participant, bar height and error bars the group mean and standard error. H/MAM: hits/misses AM noise, NT: task-irrelevant NT tones, H/MNT: hits/misses NT tones, signal absent: signal-absent trials without button press (correct rejections).

These results reveal a parallel between PDR and AC activity, both of which were evoked by detected *and* missed stimuli when task-relevant, although weaker for misses. The response to NT tones is of particular interest here, because the same stimuli did not evoke a PDR or significant activity in AC when they were not task-relevant, and were mostly not perceived by participants. This finding raises the question of how activity in AC is related to perception, if a weaker response is also observed for missed targets. One explanation could be that the presence of AC activity for miss trials depends on the decision criterion used for reporting stimulus presence and would diminish for more liberal criteria. This possibility was explored further in experiment 2, where each participant gave trial-by-trail ratings of the strength of their perception on a six-point scale. According to SDT^13^, the amplitudes of either AC activity or the PDR should show a monotonic relationship with perception rating.

### Confidence rating of near-threshold tone perception

To obtain a behavioral rating for each NT tone, including the tones rated undetected in the binary classification, the tones were now presented in short intervals of white noise (Fig. 5A). After each interval, participants rated the strength of their perception of the tones on a six-point scale. To simplify understanding and usage of the scale, the different ratings were presented to participants as levels of confidence in tone perception, ranging from high confidence in signal presence to high confidence in signal absence. Ratings 1-3 should be used for a “tone absent more likely” choice with high, medium, or low confidence, ratings 4-6 should correspondingly be used for “tone present more likely” choices. These six ratings for signal-present and signal-absent trials lead to a total of twelve possible outcomes. The ratings obtained with this scale were relatively similar to those obtained with a continuous rating of tone audibility (Fig. S3). One advantage of the 6-level confidence scale we used seems to be more consistent use of the entire range. For the analysis of the individual ratings, we expected a gradual increase of neural activity across ratings with the smallest amplitudes for rating 1 (the most confident misses) and the largest ones for ratings 6 (the most confident hits).

**Fig. 5.**
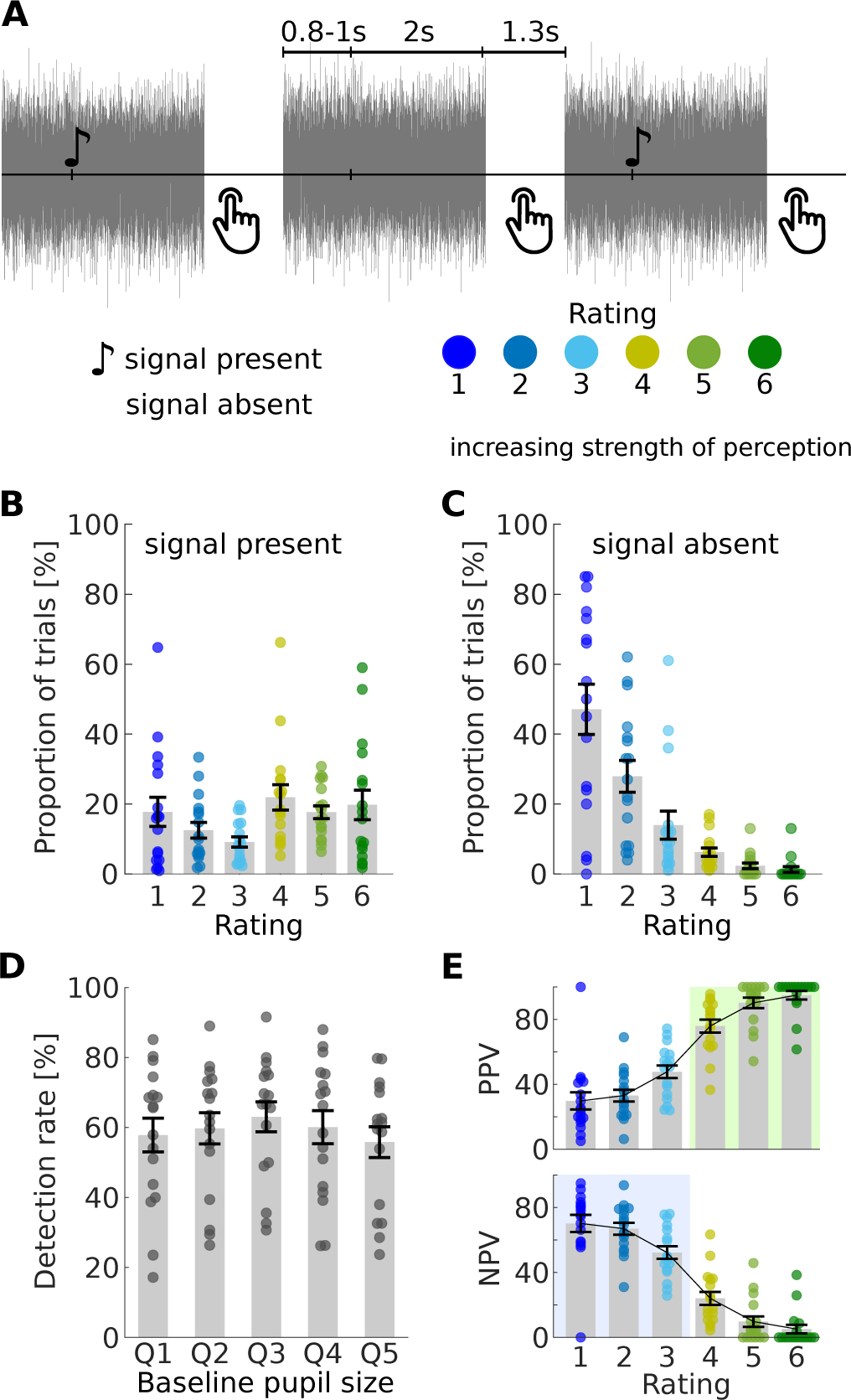
Paradigm and behavioral results of Experiment 2 (Rating task). A) Experimental paradigm: NT tones and signal-absent trials are presented in intervals of white noise. After each noise interval, the perception of a tone is rated via button press on a six-point scale. B)-E) Behavioral results. Single points represent single participants, bar height and error bars represent the group mean and standard error. B)+C)Probability for the different outcomes for signal present (B) and signal absent (C) trials. Note that there were more signal-present than signal-absent trials (Npresent=500 vs. Nabsent=100 for each participant). D) Detection rate as a function of baseline pupil size (quintiles). E) Positive (upper) and negative (lower) predictive values for each rating.

### Behavior

The relative frequency of each response is shown in Fig. 5B. In the signal-present trials, responses were spread out almost equally across ratings 4-6 and rating 1, while rating 2 and 3 were chosen less frequently. For signal-absent trials, in contrast, there was a strong bias towards low ratings (Fig. 5C). When the analysis of target trials was dichotomized in the middle of the scale, i.e. into hits vs. misses with rating 4-6 vs. 1-3, the mean hit and false-alarm rates were 59.4 ± 18.5% and 9.9 ± 9.2% (mean ± s.d.), respectively, resulting in an average decision criterion of 0.6 ± 0.4. Accordingly, the approach of offering multiple response options to lower and spread the response criterion was successful: With a criterion similar to the average of Experiment 1 (c=1.2), the cut-off would have been higher, counting only rating 5 and 6 as hits, as a dichotomization below 5 would correspond to c=1.2 ± 0.5. In other words, the range of NT miss trials in experiment 1 is now spread into ratings 1 – 4, allowing for the test of a correspondence between rating and neural response strength of supposedly unperceived trials.

When the dichotomized hit rate for combined ratings 4-6 is plotted against pre-stimulus pupil size (Fig. 5D), it increases from low to medium pupil sizes and decreases afterwards. A contrast analysis showed a significant quadratic effect (F_1,16_=15.8, p=0.001) and no significant linear effect (F_1,16_=0.4, p=0.53). This is in line with previous publications^8,22^ that report an inverted U-shaped relationship between performance (either accuracy or reaction time) and arousal. Compared to our first experiment, the noise-onset before each trial most likely increased overall arousal (i.e., pre-stimulus pupil size), enabling us to observe performance across a broader range of arousal levels.

It would be reasonable to assume that this also caused the increased hit rates compared to the first experiment. However, when only those participants who listened to NT tones with a signal-to-noise ratio of -21dB are compared (N=13 for experiments 1, N=12 for experiment 2; see Methods for details), the change in hit rate (32 ± 17% in experiment 1 vs. 55 ± 19% in experiment 2, t_23_=-3.2, p=0.0038) and criterion (1.14 ± 0.42 in experiment 1 vs. 0.56 ± 0.38 in experiment 2, t_23_=3.6, p=0.0016) is similar, but the detectability d’ does not change significantly (1.2 ± 0.6 for experiment 1 vs. 1.4 ± 0.6 for experiment 2, t_23_=0.8, p=0.42). Consequently, the increased hit rates can be attributed solely to the lower criterion. Accordingly, when only ratings 5 and 6 are combined to calculate a dichotomized hit rate with a criterion corresponding to that of the first experiment, the resulting hit rate of 37 ± 18% is also in a similar range.

Additionally, we calculated the positive and negative predictive values (PPV and NPV) for each rating, i.e. the ratio of correct “signal present” or “signal absent” responses. With labeling responses as correct or incorrect, this approach implies a threshold: Ratings above threshold are considered as correct for signal-present trials, and incorrect for signal-absent trials. Because the predictive values are calculated separately for each rating, they do not change for different thresholds, as long as they are lower (for PPV) or higher (for NPV) than the respective rating. In other words: The PPV for a response with rating 6 remains the same, whether the threshold is assumed between rating 5 and 6 or below rating 1. Therefore, the PPVs in Fig. 5E are based on an assumed threshold below rating 1, and the NPVs are based on an assumed threshold above rating 6. For any given threshold in between, only the values for ratings above/below that threshold should be considered.

The results in Fig. 5E show an increase of PPV and a decrease of NPV across the ratings (rmANOVA: F_5,80_=56.0, p<0.0001). A threshold in the middle of the scale between ratings 3 and 4 would, for example, result in three PPVs and NPVs each (PPV 4-6; NPV 1-3; Fig. 5E, colored background). For both, the effect across the three ratings is significant (F_2,32_=19.2, p<0.0001 for PPV; F_2,32_=5.0, p=0.027 for NPV). A significant effect across NPVs would not be expected in a threshold-based model of perception, as listeners would not have access to the different levels of activity on trials below threshold. Therefore, ratings on unperceived trials would be guesses and the NPV should be the same for all sub-threshold ratings^26^. Consequently, if a threshold was assumed, nevertheless, it would need to be below rating 3.

### M/EEG

Qualitatively, the M/EEG results (Fig. 6A-B, Fig. S2) were similar to those from Experiment 1, but the overall amplitude of the evoked response was lower by a factor of 2. The lower amplitudes observed in Experiment 2 could be due to the short time interval between noise and tone onsets, whereas in experiment 1, continuous noise was used, and thus the preceding auditory event was typically the previous NT tone (or AM noise). Together with the relatively low number of signal-absent trials (n=100 for each participant), SNR was too low for the analysis of correct rejections and false alarms across all ratings. Therefore, correct rejections were combined across ratings 1 – 3. False alarms (ratings 4-6) were too infrequent to be analyzed further.

**Fig. 6.**
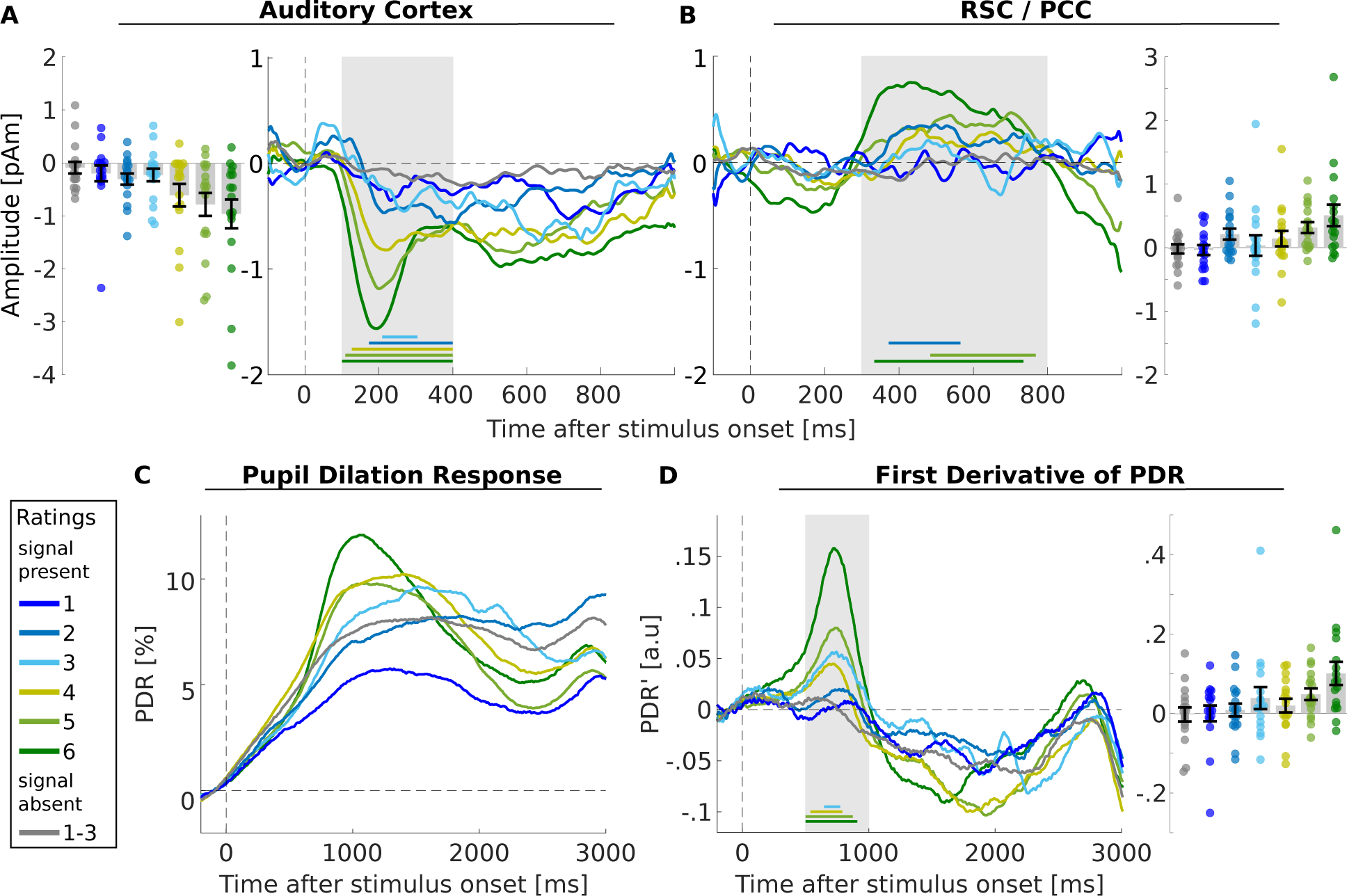
M/EEG waveforms and pupil dilation response of Experiment 2 (Rating task). Time courses of bilateral AC (A) RSC/PCC (B), PDR (C) and PDR’ (D) together with mean amplitudes in the time windows of interest (gray background in the waveform plots, AC: 100-400 ms, RSC/PCC: 300-800 ms, PDR’: 500-1000 ms; no amplitudes for the PDR, because the PDR’ was used for statistical tests). The colored horizontal bars in the waveform plots mark the time windows of significant activity (permutation cluster test). In the amplitude plots, each circle represents the respective amplitude of one participant, bar height and error bars represent the group mean and standard error.

AC source activity shows that the amplitude of the AAN for detected targets increases monotonically as a function of perceptual rating (Fig. 6A), with the mean amplitude of rating 3 being the lone exception (perhaps due to the low SNR of that condition as the least frequently chosen option). A contrast analysis confirmed a linear relationship between AC source activity and rating (linear: F_1,16_=10.7, p=0.005; quadratic: F_1,16_=0.7, p=0.43). In the permutation cluster test between 100 and 400 ms, the AAN was significantly different from zero for all conditions except rating 1. Preceding the AAN, a positive-going P1 was observed, more prominently for rating 1-3, where the AAN was smaller. A similar result has been previously observed in the context of informational masking, where a P1 was only observed for miss trials^24^, perhaps because it was obliterated by the more prominent AAN in hit trials. A P2-like response, as in the first set of experiment 1, was not observed for low ratings of experiment 1.

RSC/PCC source activity shows a prominent response for rating 6, consistent with a P3-, with smaller amplitudes and varying duration and consistency for ratings 2-5 (Fig. 6B). But only ratings 2, 5, and 6 were significant in the permutation cluster test between 300 and 800 ms. A linear amplitude effect across all ratings was significant in a contrast analysis (linear: F_1,16_=7.5, p=0.014; quadratic: F_1,16_=2.5, p=0.135).

### Pupil dilation response

The PDR for stimulus onset increases through baseline for all trials in response to the noise onset, which occurred 0.8-1 second before a target tone (if a target tone was present). The PDR peaks around 1000 ms after tone onset and increased in amplitude from rating 1 to 6, with the PDR for the smallest ratings or 1 or 2 being even smaller than that for correct-rejection trials (gray trace in Fig. 6C). To separate the PDR to targets from the PDR to noise, the first derivative of the PDR was calculated and additionally baseline-corrected from -200-0 ms prior to target-tone onset. The resulting PDR’ (Fig. 6D) confirms the linear increase of the tone-evoked PDR from rating 1 to 6 (contrast analysis, linear: F_1,16_=12.8, p=0.003; quadratic: F_1,16_=1.4, p=0.253). Despite the delayed button press, the first derivatives are similar to those of experiment 1, suggesting that the button press itself did not have a relevant impact on the early portion of the PDR that was analyzed here.

### Modeling neural activity based on the behavioral results

The relationship between response amplitude and rating is generally in line with the idea of a continuous range of neural representation being directly linked to perception. To examine this relationship in more detail, the within-participant effect of decision criterion was used to model the evoked activity based on SDT^13^. The model assumes that signal strength for signal and noise trials are normally distributed with different means and standard deviations. The mean amplitude for each decision criterion can be calculated using only the probability of each trial type, i.e. the frequency with which each response occurred. We chose the parameters of the noise distribution as μ_noise_ = 0 and σ_noise_ = 1. Following the steps described in the method section, we calculated μ_signal_ = 1.385 and σ_signal_ = 1.344. This results in a ratio of Δμ/Δσ = 4.02, which is in line with the empirical constant of Δμ/Δσ = 4 suggested by Green and Swets^13^ (This ratio describes the assumption of unequal variance, i.e. that the variance of the signal and noise distribution vary, and was confirmed across different studies). The resulting distributions for signal and noise trials and the predicted and measured amplitudes are plotted in Fig. 7A, along with a bifurcation-based model (Fig. 7B) and a threshold-based model (Fig. 7C) (see below). The predicted curve is similar to the measured activity in AC (R² = 0.85). The corresponding results for RSC/PCC and PDR can be found in the supplement (Fig. S4).

**Fig. 7.**
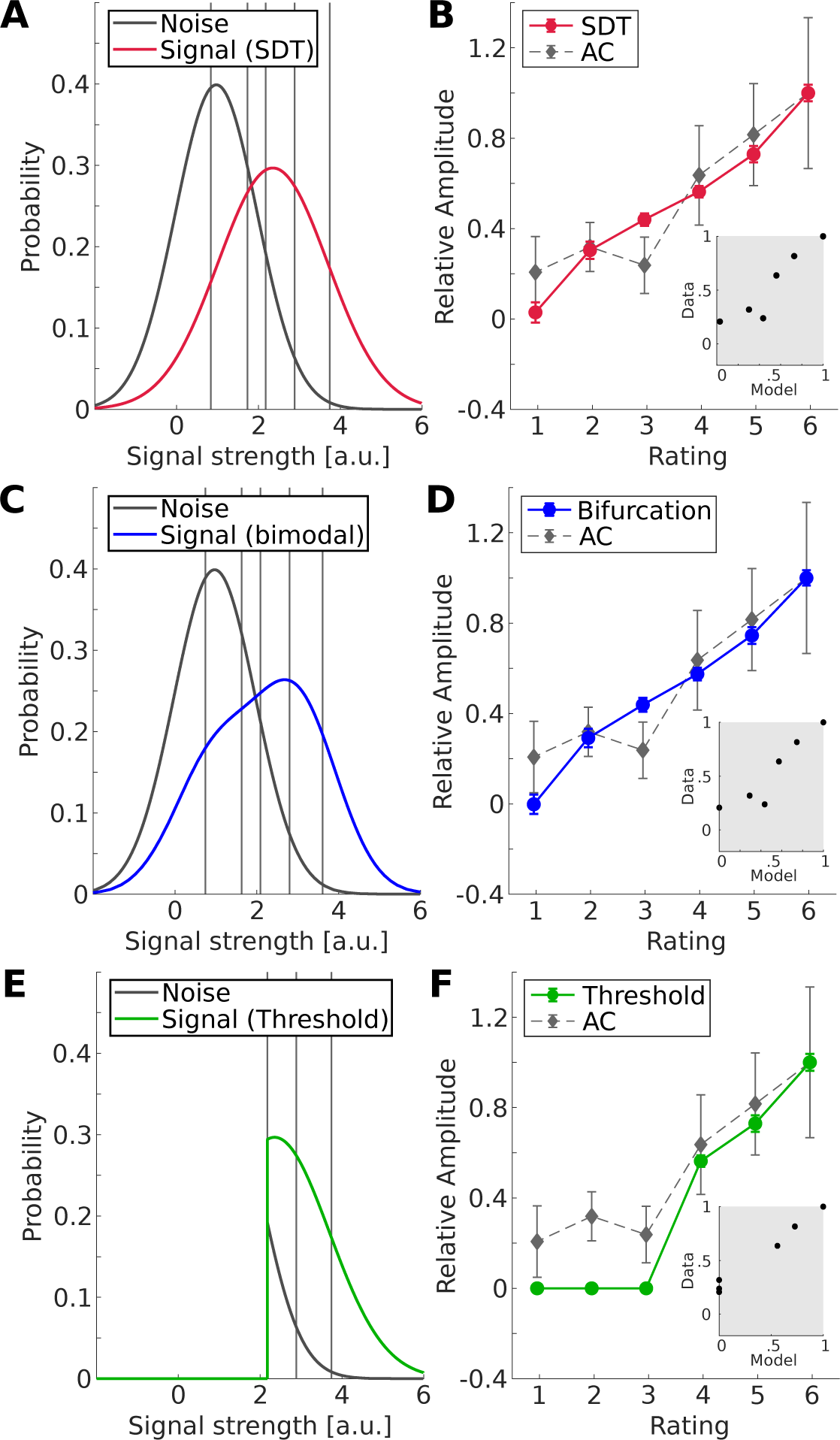
Modeling of neural activity based on behavioral results of Experiment 2 (Rating task). (A) Signal (red) and noise (black) distribution fitted with a STD-Model based on the behavioral results. Vertical lines represent the group average of the five criteria calculated based on the two distributions. Note that due to of the arbitrary choice of μnoise, the resulting criteria do not exactly follow the definition of equation (1), but are shifted by a fixed value corresponding to the intersection of the two distributions. (B) Prediction of relative amplitudes (solid line/circles) and actual amplitudes of AC activity (gray dashed line/diamonds). All data points are averages across participants, error bars represent the standard error of the mean. The inset shows the predicted vs. the measured amplitudes. (C) and (D) are the equivalent plots for a model with a bimodal signal distribution ^6^, (E) and (F) for a threshold model, which assumes no AC activity below threshold.

The results show that the early activation in AC can be modeled using a continuous, unimodal signal distribution, suggesting that the percept and neural representation of NT auditory signals are continuous rather than discrete. Next, we compared the SDT model with the recently proposed bifurcation model, where a bimodal signal distribution is used^6^. As can be seen in Fig. 7B, the difference between the classical SDT model and the bifurcation model are marginal: The respective signal distributions are highly similar (Pearson’s r=0.98, p<<0.0001) and the bimodal model provides a similar explanation of the AAN data (R² = 0.83). As another alternative, a threshold model is shown in Fig. 7C. This model assumes that sub-threshold trials do not evoke any activity at the neural level related to perception, although such activity is expected to be present in lower levels of sensory processing. The level for all sub-threshold activity is assumed to be zero, in line with the activity in noise-only trials, which was not significantly different from baseline in our data. The threshold model therefore underestimates the activity for trials below the assumed threshold (ratings 1-3), which is reflected by less variance explained by this model (R² = 0.61). Setting the threshold between 3 and 4 is arbitrary, and some threshold models have argued for higher or lower thresholds^26^. The further the threshold is shifted to the left, the more the threshold and SDT models become similar and eventually indistinguishable, as there is no way to differentiate a low-threshold model from a threshold-free model in this experiment.

## Discussion

The present data show that AC activity for ambiguous NT tones depends strongly on attention, such that no negative-going activity was observed when the tones were not task-relevant and remained mostly unperceived. When the tones were task-relevant, however, prominent activity was observed for all tones, though much stronger when detected, when a simple detection task was used in experiment 1. When the tones were again not task-relevant in run 3, half of the listeners denied having heard them, despite being informed about their presence after run 1.

This strong task and (likely) attention effect on tone detection could be considered inattentional deafness, and may in fact be stronger than effects that have been reported in previous multi-phase experiments of inattentional blindness^19^ and deafness^20^, where the d’ of the stimuli was greater than 3 and the stimuli were detected by 20%^20^ and 50%^19^ of participants before debriefing (and 100% after it).

The AAN recorded from AC in the present experiment has been proposed as a neural correlate of perceptual awareness^2,27,28^. However, one critique of this interpretation is that the AAN reflects attention rather than awareness^29^. The presence of similar, negative-going responses for undetected tones is a further argument against a close relationship with awareness, based on the assumption that a neural correlate of awareness would be observed only for hit trials (note that the AAN is often defined as difference between aware and unaware trials^2,27^). Such a strict separation of consciously perceived and unperceived tones is particularly prevalent in threshold models of perceptual awareness^30^. In such a model, the observed enhancement for miss trials would be an instance of attentional stimulus enhancement in the absence of perceptual awareness, and accordingly dissociate attention and awareness for this particular instance. Auditory cortex activation for missed trials has been demonstrated before^16,31^; what’s novel about the present results is that AC activity is enhanced, i.e. depends on attention, in task-relevant miss trials compared to a situation where the same stimuli were not task-relevant.

Alternatively, the transition between hit and miss trials can be modeled continuously, as is suggested by classical SDT^13^. In this case, the decision for hit and miss trials is based on a response criterion that can be shifted to produce more hits, but at the cost of more false alarms. While the relationship between response criterion and N1 (or AAN) amplitude has been observed before^32^., the present data extend these results and show that the AAN can be observed for much lower criteria than shown previously, i.e., well within the range of typically observed “miss” trials in binary rating experiments. Moreover, the results of this experiment are consistent with SDT, as demonstrated by the relationship between AC activity and the SDT model. A limitation of our results is that we cannot demonstrate a monotonic effect of rating on AAN amplitude in the range of criteria below 1.2. However, a significant effect of response accuracy in the range of ratings 1-3 strongly suggests that participants can dissociate these levels, further arguing for a threshold-free model.

More recently, a bifurcation model has been suggested where the dichotomy of hit and miss trials is explained by a bimodal distribution of neural activity instead of the unimodal Gaussian distribution used by SDT^6^. However, as the simulation for the single intensity, ambiguous NT tone used in our study indicates, the difference between the bimodal distribution suggested by the bifurcation model compared to unimodal distribution from classical SDT is negligible, and below the noise in our data.

Irrespective of whether classical SDT or a bifurcation model better accounts for our data, the more important question for interpretation of the AAN is whether the model assumes a threshold or not. If it does, the same restrictions apply as outlined above. If it does not, but was considered similar to SDT other than the signal distribution (as is assumed in Fig. 7), then the model is well in line with our data, but would also predict a continuous transition from perceptual awareness to unperceived trials for the NT stimuli used here. As outlined above, the dissociation between threshold and SDT models is important to interpret the AC response. In the SDT model, the presence of the AAN for miss trials does not conflict with the character of an awareness-related response. In contrast, threshold models require that an awareness response is observed only for hit trials, and our data were not well fit by such a threshold-based model^5^. Accordingly, the present data support the interpretation of a close correspondence between the AAN and conscious auditory perception under the assumption of a threshold-free, SDT model. Note that a separation of the two models, threshold versus non-threshold, would predict differences in particular on the miss side of our scale, i.e. below threshold. This distinction is particularly difficult for low-threshold models, where the difference would only be observed between very low-amplitude conditions.

While our scale was introduced to the participants by levels of confidence in the response, it can alternatively be considered a scale of perceptual strength (Fig. S3). This is of particular interest for the interpretation of the PDR’, which scaled continuously with the perceptual strength of the tones rather than decision confidence (which was highest for levels 1 and 6). In contrast, others have observed the PDR scaling with decision confidence independently of perception^33–36^. In these studies, however, the rating was secondary, and the PDR was coupled to the response rather than directly to the stimulus. As it has been shown in a multitude of cognitive tasks, how the different aspects of the PDR manifest and are affected by task instructions is likely context- and paradigm-dependent. The finding of a relationship between criterion and the PDR in visual perception^34^, on the other hand, is very similar to our finding. However, while the present study also finds a signal in auditory cortex that scales with criterion, the cited visual study did not find such a correspondence in visual cortex using fMRI. This potential coupling of PDR and processing in sensory cortex is thus the important finding of the present study, rather than the coupling of PDR with perception (or criterion), *per se*.

The next question following from this finding is what the functional coupling between AC and networks underlying the PDR could be. The response latency of LC activity supposedly driving the PDR can only be approximated based on data from a monkey model^37^: the change in pupil diameter peaked around 500 ms after microstimulation of the LC and other midbrain sites, and the delay between spontaneous LC activity and pupil dilation was in a range of 300 to 400 ms. Based on the peak latency of our PDR’ around 750 ms, LC activity in our experiment would then be expected to have a peak latency between 250 and 450 ms after stimulus onset, placing the hypothetical activity underlying the PDR in a latency range between AC and RSC/PCC. One possibility could therefore be that the AAN is related to postsynaptic activity in apical dendrites of pyramidal neurons with cell bodies in layer 5B^38^, and specifically to the subgroup that projects to subcortical nuclei. Calcium currents in apical dendrites of such layer-5B neurons in mouse somatosensory cortex have been shown to be correlated with the detection of near-threshold stimuli, in particular the subpopulation projecting to subcortical targets^12^. Less likely, the AAN may be driven by feedback projections emanating from subcortical centers. For example, late activity in the somatosensory cortex of mice is observed for salient stimuli, and for non-salient stimuli when combined with phasic, optogenetic stimulation of LC neurons^39^. It is generally possible that LC or other^10,40,41^ subcortical networks such as superior colliculus or the pulvinar also play a role for sustained attention that is required for the enhancement of NT tones in AC. Subcortical and cortical attention networks are coupled, e.g. anterior cingulate cortex and LC, or the superior temporal cortex and SC^18^, and it is clear that sustained attention has a strong cortical component^42^. Previous fMRI data have shown that pre-stimulus activity in anterior cingulate cortex and insula is higher in trials were near-threshold tones are detected^16^. While a weak link between baseline PDR and detection is also observed in the present data, this appears to be weaker than observed in the cortical attention system^16^, suggesting that the PDR does not simply reflect the cortical attention systems. Nevertheless, the observation of pre-stimulus and stimulus-evoked correlations between PDR and perception suggest that subcortical processes may also play a role for response modulation in auditory (sensory) cortex required for perceptual awareness of NT tones, in concert with cortical attention systems.

In summary, these data are consistent with a threshold-free SDT model where the neural correlate of the perceptual strength is attention-related activity in AC. Based on such a model, the AAN would be attention dependent and present in miss trials but could still be a neural correlate of perceptual awareness, as suggested previously^27,28,43^. It remains to be seen if this model can be transferred to more salient^31^ or complex^44,45^ stimuli. We think that the strong correlation of perception and a fully attention-dependent response in the present data is due to the weak signal strength and very low level of evoked responses that are less modulated by attention. The role that attention plays for perceptual awareness of salient, unmasked stimuli cannot be inferred from the present data and will require independent evidence.

## Methods

### Participants

14 participants were included in experiment 1 (6 male, 21-32 years of age, mean 25.4) and 17 in experiment 2 (9 male, 19-35 years of age, mean 24.4). One participant took part in both experiments. None of the participants provided a history of hearing disorder and all reported normal hearing. All participants provided written informed consent. The study was conducted in accordance with the Declaration of Helsinki (2013 revision) and approved by the ethics committee of the Medical Faculty at Heidelberg University.

### Stimuli and Procedure

#### Experiment 1: Selective attention task

Experiment 1 comprised three runs with different task instructions. In each run, the same auditory stimulation was presented to the listeners. It consisted of continuous white noise combined with two different types of stimuli:

1) Pure tones with a frequency of 1000 Hz and a duration of 100 ms were presented at a near-threshold level of -21 dB relative to the noise, aiming for a detection rate of 50%. This was increased to -19 dB for one participant.

2) Transient amplitude modulations of the noise with a duration of 100 ms and a modulation frequency of 50 Hz. The modulation depth was usually set to 25%, which yielded a detection rate >90% in the group of listeners used to adjust the stimuli; it was increased to 35% for four participants, because their detection performance was not reliable during the example sets.

Additionally, catch trials were included by presenting a subset of tones at a level of -100dB, equivalent to signal-absent trials. Each run comprised 150 trials with near-threshold (NT) tones, 50 signal-absent trials, and 50 trials with amplitude modulated (AM) noise. The order was randomized and the stimulus-onset asynchrony was jittered between 3 and 5 s, leading to a total duration of 17 min for each run.

Before the first run, participants were introduced to the AM noise by explaining the sound and playing short examples. If the participant was not able to reliably detect the AM noise at a modulation depth of 25%, the depth was increased. At this stage, participants were not informed about the presence of the NT tones. Afterwards, the participant was instructed to press a button as soon as an AM noise was detected, and the first run was started. Before the second run, participants performed a single-interval adaptive procedure^21^. During this procedure, short intervals of noise with or without a tone were presented. After each interval, participants indicated via button press if they heard a tone and the SNR was adjusted accordingly (lowered after detections, raised after missed tones). After eleven response reversals (from detections to a miss or vice versa), the SNRs from the last seven reversals are averaged to yield an estimate of the 50 % detection threshold. However, in pilot measurements the use of this threshold estimate resulted in a substantially different performance during the actual, non-adaptive testing. Therefore the SNR was not adjusted to this threshold estimate, but the adaptive test was rather used to familiarize participants with the stimuli. A possible reason for the discrepancy between adaptive and non-adaptive tasks is that the adaptive task would have required several repetitions to provide a stable level, which would have excessively prolonged the experiment. The instruction for the second run was to ignore the AM noise and instead detect the NT tones via button press. The performance was observed in real time, and if a participant was not able to detect any tones in the first minutes, the run was stopped and started over with a higher SNR. This was the case for only one participant (new SNR: -19 dB). For the third run, participants were instructed to repeat the task from the first run, i.e. to detect AM noise. Participants were not explicitly informed about the presence of the tones.

#### Experiment 2: Rating task

In Experiment 2, only the NT tones were used as target stimuli. In contrast to the first experiment, the white-noise masker was not continuous but split into intervals with an average duration of 3s: In each interval, the tone was presented after 800-1200 ms and the noise afterwards lasted for another 1.9 s. Tones were the same as in Experiment 1 and presented at either -21 dB (signal-present trials) or -100 dB (signal-absent trials). For participants who did not detect any NT tones in the early portion of the experiment, the SNR was increased and the experiment restarted. This was the case for five participants where the SNR was increased to -19 dB (three), -19.5 dB (one) and -20 dB (one). Participants had to rate their perception of the tone on a six-point scale ranging from “tone very certainly absent” (1) to “tone very certainly absent” (6). To do so, they had three buttons for the right hand to indicate perceived tones (mapping to “very certainly present, somewhat certainly present, and possibly present but unsure”) and three buttons on the left indicating that no tone was perceived (mapping to “very certainly absent, somewhat certainly absent, and possibly absent but unsure”). The briefing was in German, so the original assignment of the keys was “ganz sicher, ziemlich sicher, unsicher aber eher (nicht) vorhanden” for the signal present (absent) ratings. A six-point rating was chosen to enable to comparison of evoked responses with different simultaneously held decision criteria. The certainty of perception was used to differentiate between different the different ratings, because it is an intuitively easy to use scale that does not require extensive explanation or practice. A comparison of this scale and a continuous audibility rating was carried out in a supplementary experiment (Fig. S3).

Instructions were to decide fast without overthinking, which was enforced by a short, silent inter-trial interval (1 second for the first participant, and 1.3 seconds for all subsequent participants). The experiment was split in two parts to reduce fatigue. Each part consisted of 250 signal-present trials and 50 signal-absent trials and lasted approximately 20 min. The same adaptive procedure as in Experiment 1 was used before the first run to make the participants familiar with the stimuli.

All stimuli were created in MATLAB (MathWorks, Natwick, MA, USA) with 32 bit resolution and a sampling rate of 48000 Hz. An in-house, MATLAB-based software and PsychToolbox 3^46^ were used for presentation, together with a programmable attenuator (PA5, Tucker-Davis Technologies (TDT), Alachua, FL, USA), headphone amplifier (HB7, TDT), and ER-3 earphones with custom 97 cm long tubes (Etymotic Research, Elk Grove Village, IL, USA).

### Data acquisition

#### MEG and EEG

MEG was recorded with a 122-channel whole-head system (MEGIN Oy, Helsinki, Finland). In the first experiment, EEG was recorded in parallel with a 63-channel equidistant cap (Easycap, Herrsching, Germany) where the electrodes extend far down from the low cheeks until below the inion. Additionally, one ECG channel was recorded. In the second experiment, 32 EEG channels arranged according to the standard 10-20 system were recorded. MEG and EEG data were recorded with a sampling rate of 1000 Hz and low-pass filtered with 330 Hz. Prior to data acquisition, the nasion, left and right pre-auricular points, all electrode positions, and additional head shape points were digitized using a Polhemus, 3D Space, Isotrack2 system. The participant’s head position in the MEG was measured before each run.

#### Pupil size

An Eyelink 1000 Plus eyetracker (SR Research, Ottawa, Canada) was used for monocular tracking of the right eye. The sampling rate was 1000 Hz and pupil area as well as gaze position were recorded. Before each run, the eyetracker was calibrated and participants were asked to fixate a cross presented in the middle of the screen during the runs. PsychToolbox 3^46^ was used to present the eyetracking elements (calibration and fixation screen) and to synchronize recording with the stimulation.

#### Structural MRI

For source analysis of the MEG/EEG data, structural MRIs of all participants were obtained. Two full brain scans with T1 and T2 contrast were acquired using a 3-T Siemens Magnetom Trio (Siemens Medical Systems, Erlangen, Germany). For the T1 MPRAGE images, the field-of-view consisted of 256×256×192 1 mm^3^ voxels recorded with TR=1570 ms, TE=2.67 ms, TI=900 ms and a flip angle of 9°. The T2-weighted FLAIR sequence covered 192×256×25 voxels with size 0.9×0.9×5.5 mm³ and used the following settings: TR=8500 ms, TE=135 ms, TI=2400 ms and a flip angle of 170°.

### Data analysis

#### Behavioral data

In both experiments, button presses were considered as valid when they occurred within 1.3s after the stimulus (Experiment 1) or the noise offset (Experiment 2). In experiment 1, this would result in a hit (H) or false alarm (FA) if the preceding stimulus was a target or distractor, respectively. Targets or distractors without a button press in this time window were considered as miss (M) or correct rejection (CR), respectively. Button presses outside this time window were counted as false alarm.

The six ratings used in Experiment 2resulted in twelve possible outcomes for signal-present and signal-absent trials. A dichtomized hit and false-alarm rate was calculated by combining the trials across ratings 4 to 6.

For both experiments, the decision criteria were calculated for each participant as the negative average of the z-scored hit and false-alarm rates^47^.

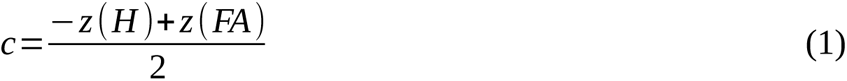

Furthermore, in both experiments, the dependence of performance on arousal was analyzed by grouping the epochs of each participant into quintiles according to the respective baseline pupil size (as a proxy for arousal), which was determined as the mean value in the 200 ms before stimulus onset. Hit rates were calculated for each quintile and averaged across participants. In Experiment 2, this was first done with the previously used dichotomized hit rate. But a post-hoc control for a split between ratings 1-5 (misses) and 6 (hits), i.e. according to a criterion comparable to Experiment 1, confirmed the effect.

The detection rates of run 2 in the first experiment were analyzed for an influence of time, as performance was lower than expected. For this aim, trials were kept in chronological order and split into five equally large parts. The detection rates for each of these time periods were calculated and averaged across participants.

In the second experiment, the positive and negative predictive values (PPV and NPV) for each rating were calculated. These values correspond to the response accuracy for signal-present and signal-absent ratings^26^. and the calculation assumes a threshold, i.e. a bimodal allocation of ratings as “signal present” or “signal absent”. In this framework, responses can be labeled correct and incorrect, depending on whether it was a signal-present or a signal-absent trial. The PPV, or “signal present” accuracy, was calculated as rate of correct signal-present responses with a specific rating (hit rate HR) divided by the sum of HR and false alarm rate (FAR): A_present,r_ = HR_r_/(HR_r_+FAR_r_) for rating r. The same was done for the NPV, or “signal absent” accuracy, using the CR and M rates (CRR, MR): A_absent,r_ = CRR_r_/(CRR_r_+MR_r_). Each rate is calculated as the number of responses with the respective rating divided by the total number of signal-present or signal-absent trials, respectively. To receive a predictive value for all possible thresholds, we calculated the PPV for each rating assuming a threshold below that criterion, i.e. assuming each response with this rating was a “signal present” rating. Consequently, all responses to signal-present trials with that rating were considered as correct (H), and to signal-absent trials as incorrect (FA). For the NPV, we correspondingly assumed a threshold above each rating, leading to correct responses (CR) for signal-absent trials, and incorrect responses (misses, M) for signal-present trials. Consequently, for the whole set of predictive values of a certain threshold, one would consider the NPVs of ratings below the threshold, together with the PPVs of ratings above the threshold. For example, an assumed threshold between rating 3 and 4 would result in NPVs for ratings 1-3, and PPVs for ratings 4-6.

#### MEG and EEG

MNE-Python v0.18 and v0.20^48^ was used together with Freesurfer v6.0^49–51^ for preprocessing and source analysis of MEG and EEG data. Raw data were band-pass filtered between 0.05 Hz and 40 Hz. External artifacts (for example, streetcars) were corrected with signal-space projection (SSP^52^). This method projects the signal to a subspace orthogonal to the noise signal, thereby reducing the degrees of freedom. The projections were calculated on the low-frequency part of the data (0.05 – 0.5 Hz) which covers the majority of our environmental artifacts. For each data set, six projections were calculated, but some were discarded when they did not clearly improve data quality. On average, 4.4 projections were used (average across participants of both experiments, range: 2-6). Biological artifacts (ECG and EOG) were corrected with independent component analysis (ICA), where the signal is decomposed in statistically independent components, and those components that include artifacts can be removed^48^. ICA was calculated and applied separately for MEG and EEG channels. The ECG channel used in experiment 1 was used as a reference for the cardiac artifacts, the EEG channels on the forehead for the ocular artifacts. After artifact correction, epochs with remaining, excessive noise were determined with the python package autoreject^53^ and excluded from further analysis. The average rejection thresholds were 4.3×10^-10^T/cm (SD: 3.7×10^-10^T/cm) for MEG channels and 1.7×10^-4^V (SD: 2.2×10^-4^V) for EEG channels. On average, 8% of the epochs in the first experiment and 23% of the epochs in the second experiment had to be excluded.

For source analysis, the cortical surfaces were reconstructed from the structural MRI with FreeSurfer, and individual source estimates were morphed to a brain template (fsaverage from FreeSurfer^51^). Source estimates were calculated with minimum norm estimates (MNE) and dynamic statistical parametric mapping (dSPM^54^ where the MNE signal is normalized according to the signal variability at each source location. The noise covariance matrices used for both methods were calculated on the 100 ms pre-stimulus activity. The dSPM source estimates were used to create maps of the brain activity. These maps were averaged across participants to show areas with significant activity. The medial wall (“unknown” label in the Destrieux Atlas ^55^) was excluded from the group average plots, as it likely does not include relevant brain activity but rather spread from other sources^56^. The MNE source estimates were used for the region-of-interest (ROI) analysis, because it allows for a direct comparison of the response strengths, whereas the dSPM amplitudes depend on the number of averaged trials. The crown of Heschl’s Gyrus was chosen as the expected main source of activity in auditory cortex (AC), and an area covering the posterior cingulate cortex (PCC) and the retrosplenial cortex (RSC) was chosen as the expected source of the decision-related P3 component^25^. From these two areas, waveforms were extracted by averaging the activity across all vertices in the corresponding label with MNE-Python’s “mean-flip” method, where polarities are corrected depending on the respective source orientation. The extracted waveforms were subsequently averaged across participants. For the first experiment, because tasks differed across runs, each run was analyzed separately; in the second experiment, both runs were combined.

#### Pupil size

The eyetracking data were imported to MATLAB using the edfmex toolbox provided by SR Research. Further preprocessing was done using custom scripts: Blinks were corrected with linear interpolation from 100 ms before to 100 ms after the blink. Epochs where the amount of interpolated data exceeded 20% were excluded from analysis. This affected 18% of the epochs in Experiment 1 and 28% of the epochs in Experiment 2. The data were low-pass filtered with a cut-off frequency of 4 Hz and split into epochs from 500 ms before to 3000 ms after stimulus onset. A divisive baseline correction was applied to each epoch by dividing all values by the mean pupil size in the 200 ms before stimulus onset. This transforms the pupil dilation response (PDR) into a change signal relative to the size during the individual baseline period. For statistical analysis, the first derivative, i.e. the temporal change of the pupil size was used, because is better suited to capture small differences in the initial event-related dilation of the pupil (the rising part of the PDR curve). It has been further suggested that this initial increase in pupil size tracks the fast, norepinephrine-driven changes in pupil size, as opposed to the more sustained dilation driven by acetylcholin^57^. The derivative (PDR’) was calculated based on the epochs without baseline correction, as the difference between one sample to the next. The result was smoothed with a moving mean across 200 samples, corresponding to a non-casual 5Hz low-pass filter. In the second experiment, the noise onset strongly influenced the pupil measurement, resulting in a strong increase in pupil size across all conditions including signal-absent trials. To remove this distortion, we performed an additional baseline correction of the derivative by subtracting the mean value during the baseline period.

#### Statistical analysis

Various effects on the behavioral performance were tested with repeated measures ANOVAs (rmANOVA): The change of detection rate across time in experiment 1, the dependence of performance on the baseline pupil size in both experiments (with a linear and quadratic contrast analysis), and the influence of rating on response accuracy in the second experiment. These analyses were performed with the statistics software SAS (SAS Institute Inc., Cary, NC, USA), sphericity correction was applied with the Greenhouse-Geisser method. The decision criteria for the different runs in experiment 1 were compared using a two-sample t-test in MATLAB.

The M/EEG waveforms and the first derivative of the PDRs were analyzed with a temporal permutation cluster test^23^ using MNE-Python and the python packages SciPy^54^ and NumPy^55^. The cluster forming threshold was set to 0.05 and a cluster was considered significant with a p-value of p < 0.05. First, we used this to test for differences between both hemispheres. Since no relevant differences were found, the averages across both hemispheres were used for the final analysis to enhance the signal-to-noise ratio. Ideally, each waveform should be tested against some control condition like the signal-absent trials, or more specifically the correct rejections.

However, because of the low number of signal-absent trials, this signal is noisy and could potentially cause an increased number of type-II errors when used as reference. To circumvent this problem, we created additional signal-absent “trials” in the M/EEG data sets from 1.9 to 3s after each stimulus onset (2s after the original stimulus onset will correspond to t=0s in the waveform plots). As the interval between two stimulus onsets was at least 3s, this time window should contain little to no event-related activity and is therefore suitable as a reference. Combining these “artificial” signal-absent trials with the original ones resulted in up to 300 trials per run (potentially reduced by exclusion due to artifacts).

Unfortunately, this approach was neither possible for the pupil data, which returns to baseline much slower, nor for the second experiment, where noise on- and offsets leave no suitable baseline periods. Consequently, we tested all M/EEG waveforms of the first experiment against signal-absent trials, but the first derivative of the PDR waveforms and all M/EEG waveforms of the second experiment against 0. Different time-windows were chosen for each ROI or modality: t_AC_=100-400ms, t_RSC/PCC_=300-800ms and t_Pupil_=500-1000ms. In the first experiment, the AC waveforms were additionally compared across runs with pairwise comparisons (run 1 vs. 2, 2 vs. 3, and 1 vs. 3) for both, AM noise and NT tones, respectively.

The amplitudes for each condition were calculated as averages within the corresponding time windows. To investigate if and how the amplitudes and peak latencies changed across ratings in Experiment 2, linear and quadratic contrast analyses were conducted.

### Modeling of neural activity based on behavioral results

To test whether the strength of perception, measured as the confidence rating, is directly related to the strength of neural activity, we modeled the neural activity solely based on the behavioral results, using signal detection theory (SDT^13^). SDT postulates that there are two different probability distributions of neural activity for signal and non-signal trials. The signal distribution itself is continuous and the differentiation between “perceived” and “unperceived” or between different confidence ratings is an arbitrary threshold (i.e., the decision criterion) set by the listener. Both distributions (signal and noise) are gaussian:

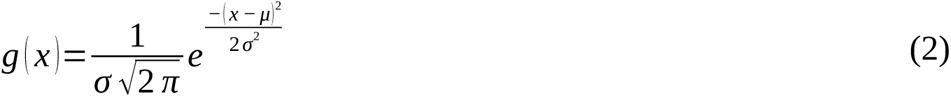

When using a six-point rating scale like ours, participants have to maintain 5 decision criteria at the same time^13^. Criterion 1 would be the most liberal, where all targets with a rating between 2 and 5 are considered as hits, while targets with rating 1 count as misses. For each more conservative criterion, one rating is removed from the “Hit” category, so that the most conservative criterion 5 includes only targets with rating 6, while all other ratings are considered as misses. Correspondingly, the same can be done for the signal-absent trials (i.e., the noise distribution) with false alarms and correct rejections.

To model the average strength of neural activity in each criterion, the distributions and the positions of the criteria are needed. The parameters μ (mean) and σ (standard deviation) of both distributions are tightly linked to each other^13^, but the absolute values are not relevant for the calculation of the criteria. Therefore, we arbitrarily set μ_noise_ = 0 and σ_noise_ = 1. With this, it is possible the criteria based on the noise distribution were calculated: The area under the curve to the right of the criterion corresponds to the probability of a trial type. For example, for criterion 1 (the most liberal), the area under the noise distribution corresponds to the probability of signal-absent trials with ratings 2-6 (the area under the signal distribution would correspond to target trials with the respective ratings). To calculate the criteria, the following equation has to be solved for c_i_ (i=1-5 for the five criteria, p_i_: probability of the corresponding events):

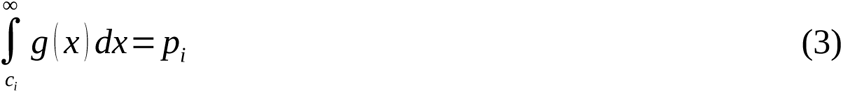

Note that due to of the arbitrary choice of μ_noise_, the resulting criteria do not exactly follow the definition of equation (1), but are shifted by a fixed value corresponding to the intersection of the two distributions. To do the same for the signal distribution, μ_signal_ and σ_signal_ are needed. The ratio σ_noise_/σ_signal_ can be calculated from the ROC curve using hit and false-alarm rates for each criterion^13^. To find μ_signal_, we calculated the five criteria for the noise distribution and for the signal distribution with different values of μ_signal_. We then chose the value of μ_signal_, where the differences between the criteria for signal and noise distribution are minimal (theoretically, they should be equal). These calculations were done for the grand average data, i.e., the probabilities for each trial type averaged across participants, to give more robust estimates of the parameters.

Based on the resulting distributions, the criteria can be calculated for each participant individually. Now the criteria should be based on both, the signal and the noise distribution. As stated above, due to the noise in real data, the above equation will most likely not result in the same c_i_ for both distributions. Therefore, the optimal choice is the c_i_ that minimizes this sum:

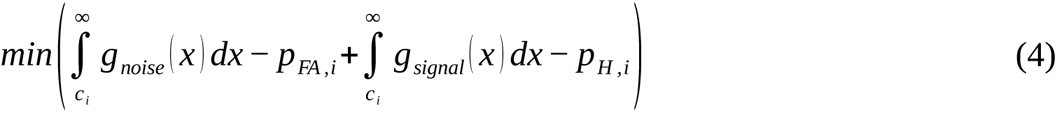

Subsequently, the mean amplitudes of each rating can be calculated using the following formula with the respective distributions *g_signal_*or *g_noise_* (i=1-6 as limits for the 6 ratings, with c_1-5_ as calculated from Eq. 3 and c_0_ /c_6_ =+/-∞):

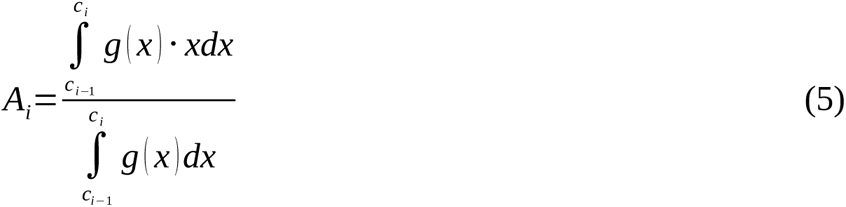

After calculating A_1_ for noise trials, i.e. the signal-absent trials with the lowest ratings, we shifted the distributions and criteria by its mean across participants, to bring the mean of the lowest rating to 0 and thus avoid negative values. Subsequently, all amplitudes were calculated. Note that this shift can influence the comparison of the model with the data or alternative models, but it is in line with the measured amplitudes for signal-absent trials.

For comparison, we constructed two additional models which consider dichotomous perception: First, a model based on a recent study^6^ where the signal distribution is bimodal and modeled as the weighted sum of two gaussians with different means but the same standard deviation for the noise distribution and both modes of the signal distribution (for more details see ^6^). We chose the weight β=0.59 according to our average dichotomized detection rate (ratings 1-3 as missed, 4-6 as detected targets), and μ_noise_ = 0 as in the previous model. From here, the procedure for calculating μ_signal_, c_i_, and A_i_ is the same as described above. Repetitions of the analysis with other weights, according to detection rates for different splits (e.g. ratings 1-5 vs. rating 6) yielded similar results. Second, a threshold model, where no activity is evoked below a fixed perceptual threshold. For this model, distributions and criteria were calculated according to SDT as described above. Subsequently, the amplitudes of the presumed sub-threshold ratings 1-3 were set to zero. Finally, to compare the resulting amplitudes with the measured AC activity (mean values in respective time windows, see above), all data sets were divided by A_H1_ for normalization before averaging across participants. We calculated the coefficient of determination R² for the average data to analyze how much of the variance in the measured data set can be explained by all of the models. The same analysis for the RSC/PCC and PDR’ amplitudes can be found in the supplement (Fig. S4).

## Author contributions

Conceptualization: AG, LD, AD

Methodology: LD, AG, AD

Investigation: LD

Formal Analysis: LD

Visualization: LD

Validation: AG, AD

Supervision: AG

Writing—original draft: LD, AG

Writing—review & editing: AG, AD

Funding acquisition: AG

Project administration: AG

