## Supplemental information for "Perceptual awareness of near-threshold tones scales gradually with auditory cortex activity and pupil dilation"

**Table S1. Results of temporal permutation cluster test.** The time courses were tested in different windows: 100-400ms in the AC, 300-800ms in the RSC/PCC and 500-1000ms in the PDR'. Significant clusters (p<0.05) are marked with an asterisk, dashes indicate, that no cluster was found. In some conditions, two separate clusters were found in the respective time window.

| | | AC | | | RSC-PCC | | | PDR' | | |
|---|---|---|---|---|---|---|---|---|---|---|
| | | *t [ms]* | *p* | | *t [ms]* | *p* | | *t [ms]* | *p* | |
| **Experiment 1** | | | | | | | | | | |
| Run 1 | detected AM | 100-230 | 0.0001 | * | 331-776 | 0.0002 | * | 500-1000 | 0.0004 | * |
| | missed AM | 125-169 | 0.0614 | | - | 0.0911 | | 629-977 | 0.0006 | * |
| | | 174-193 | 0.1069 | | - | 0.0706 | | | | |
| | NT | - | - | | - | 0.1782 | | 563-584 | 0.1490 | |
| | | | | | | | | 607-619 | 0.1730 | |
| | Catch | - | - | | - | - | | - | - | |
| Run 2 | AM | 100-219 | 0.0001 | * | - | - | | 577-737 | 0.0556 | |
| | | 311-400 | 0.0353 | * | | | | | | |
| | detected NT | 102-400 | 0.00002 | * | 395-745 | 0.0002 | * | 521-989 | 0.0002 | * |
| | missed NT | 104-400 | 0.0003 | * | - | - | | 625-1000 | 0.0016 | * |
| | Catch | - | - | | - | - | | - | - | |
| Run 3 | detected AM | 129-192 | 0.0389 | * | 335-770 | 0.00002 | * | 500-995 | 0.0002 | * |
| | missed AM | - | - | | 501-526 | 0.2158 | | 577-753 | 0.0262 | * |
| | | | | | 579-599 | 0.2323 | | 883-893 | 0.1646 | |
| | | | | | 618-682 | 0.0331 | * | 913-1000 | 0.0700 | |
| | | | | | 692-707 | 0.2702 | | | | |
| | NT | - | - | | - | - | | 647-790 | 0.0620 | |
| | Catch | - | - | | - | - | | - | - | |
| **Experiment 2** | | | | | | | | | | |
| Catch | (ratings 1-3) | - | - | | - | - | | - | - | |
| Rating | 1 | 210-305 | 0.0233 | * | 736-753 | 0.2499 | | 646-775 | 0.0499 | * |
| | 2 | 174-400 | 0.0005 | * | 374-567 | 0.0120 | * | - | - | |
| | | | | | 622-679 | 0.1177 | | | | |
| | | | | | 684-710 | 0.2483 | | | | |
| | 3 | 220-251 | 0.1116 | | - | - | | - | - | |
| | 4 | 127-400 | 0.0014 | * | 448-468 | 0.2264 | | 540-790 | 0.0310 | * |
| | | | | | 645-677 | 0.1832 | | | | |
| | | | | | 770-787 | 0.2387 | | | | |
| | 5 | 110-400 | 0.0003 | * | 406-468 | 0.1214 | * | 500-874 | 0.0002 | * |
| | | | | | 486-770 | 0.0019 | | | | |
| | 6 | 100-400 | 0.0001 | * | 435-737 | 0.0004 | * | 500-909 | 0.0003 | * |

**Table S2**. **Results of temporal permutation cluster test for Figure S2**. The time courses were tested against 0 in different windows: 100-400ms in the AC, 300-800ms in the RSC/PCC and 500-1000ms in the PDR'. Significant clusters ($p<0.05$) are marked with an asterisk, dashes indicate, that no cluster was found. In some conditions, separate clusters were found in the respective time window.

| | | **AC** | | | **RSC-PCC** | | | **PDR'** | | |
|---|---|---|---|---|---|---|---|---|---|---|
| | | *t [ms]* | *p* | | *t [ms]* | *p* | | *t [ms]* | *p* | |
| **Group A: NT perceived in run 3** | | | | | | | | | | |
| Run 1 | NT | - | - | | 598-692 | 0.0625 | | - | - | |
| Run 2 | detected NT | 106-400 | 0.0156 | * | 421-702 | 0.0156 | * | 536-991 | 0.0156 | * |
| | undetected NT | 129-400 | 0.0156 | * | - | - | | 632-1000 | 0.0156 | * |
| Run 3 | NT | - | - | | 447-488 | 0.1719 | | 673-814 | 0.0859 | |
| | | | | | 565-749 | 0.0313 | * | | | |
| **Group B: NT not perceived in run 3** | | | | | | | | | | |
| Run 1 | NT | - | - | | - | - | | 519-530 | 0.2266 | |
| | | | | | | | | 547-593 | 0.1406 | |
| | | | | | | | | 612-629 | 0.1875 | |
| Run 2 | detected NT | 178-380 | 0.0391 | * | 532-679 | 0.0234 | * | 511-521 | 0.2031 | |
| | | | | | 768-786 | 0.2344 | | 553-917 | 0.0156 | * |
| | undetected NT | 164-238 | 0.0703 | | - | - | | 854-868 | 0.2031 | |
| | | 359-400 | 0.1016 | | | | | | | |
| Run 3 | NT | - | - | | - | - | | - | - | |

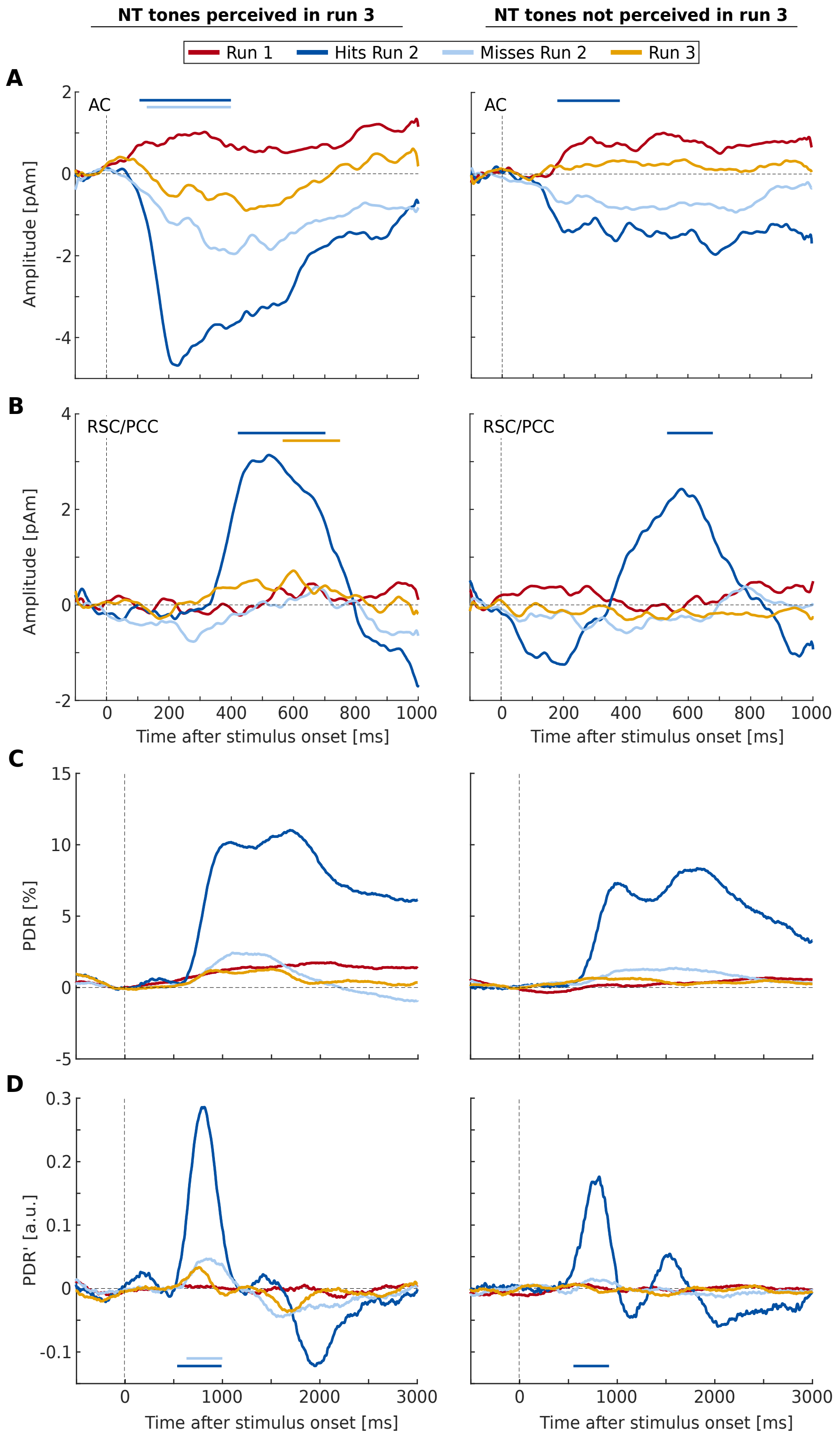


**Fig. S1** M/EEG and Pupil waveforms for NT tones in Experiment 1 for the participants who perceived tones in the last run (left column) and those who did not (right column). Time courses for the NT tones in all runs are combined in one graph each for the AC (A), RSC/PCC (B), PDR (C), and first derivative of PDR (D). The colored horizontal bars in the waveform plots mark the time windows of significant activity in the time window of interest (permutation cluster test; AC: 100-400 ms, RSC/PCC: 300-800 ms, PDR’: 500-1000 ms, no test for PDR).

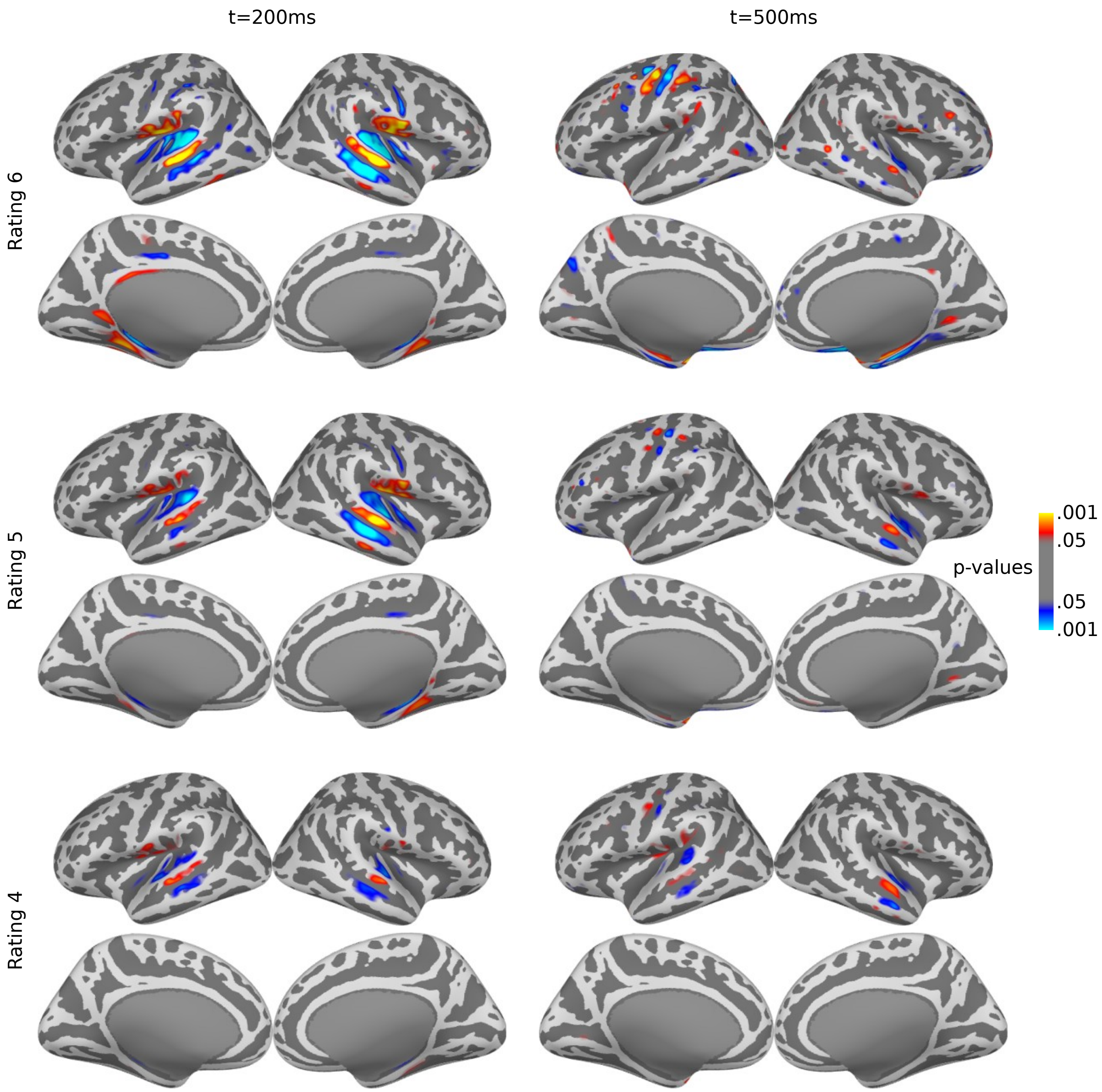


**Fig. S2. dSPM maps of Experiment 2.** Maps are only shown for rating 4-6, because there was no significant activation for the other conditions. Compared to Experiment 1, the patterns are similar, but the amplitude is greatly reduced, especially in the RSC/PCC region. Likely reasons are the trial based structure with preceding noise onset and delayed button press, and the lower number of EEG channels, since the P3b is much more prominent in EEG than MEG.

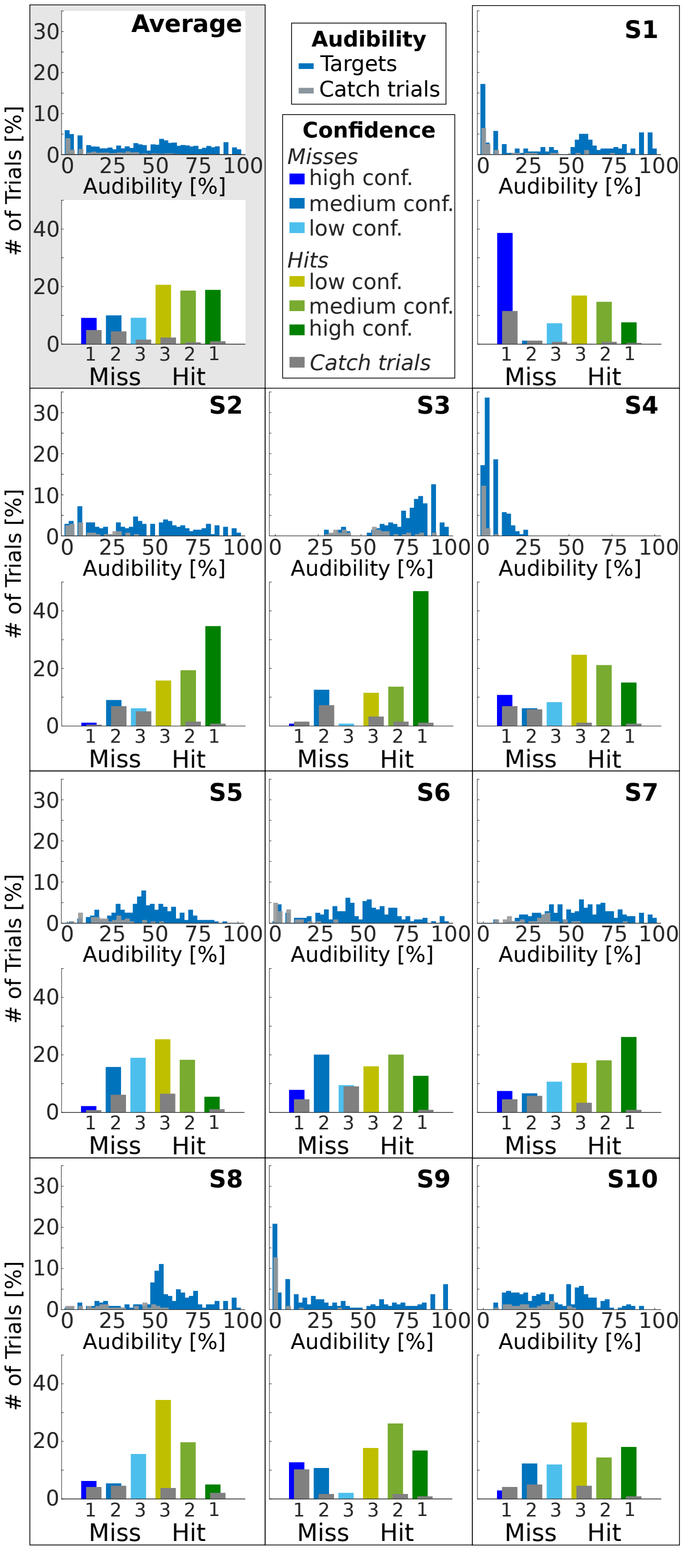


**Fig. S3 Rating scale comparison.** In an additional psychoacoustic experiment, ten participants performed two runs of tone-in-noise detection. The paradigm as in Experiment 2 was used, only the response scales differed: In the first run, participants were asked to rate the audibility of the presented tone on a scale from 0 to 100%. In the second run, they used the same confidence rating scale as in Experiment 2. In both runs, 240 targets and 40 signal-absent trials were presented to the first five participants. For the remaining five participants, the number was reduced to 210 and 35 to reduce the total duration of the experiment. In each panel, the results of the audibility rating are shown in the upper graph, while the lower graph shows the confidence ratings. The upper left panel shows the group averages for both ratings, the remaining panels are single participants data.

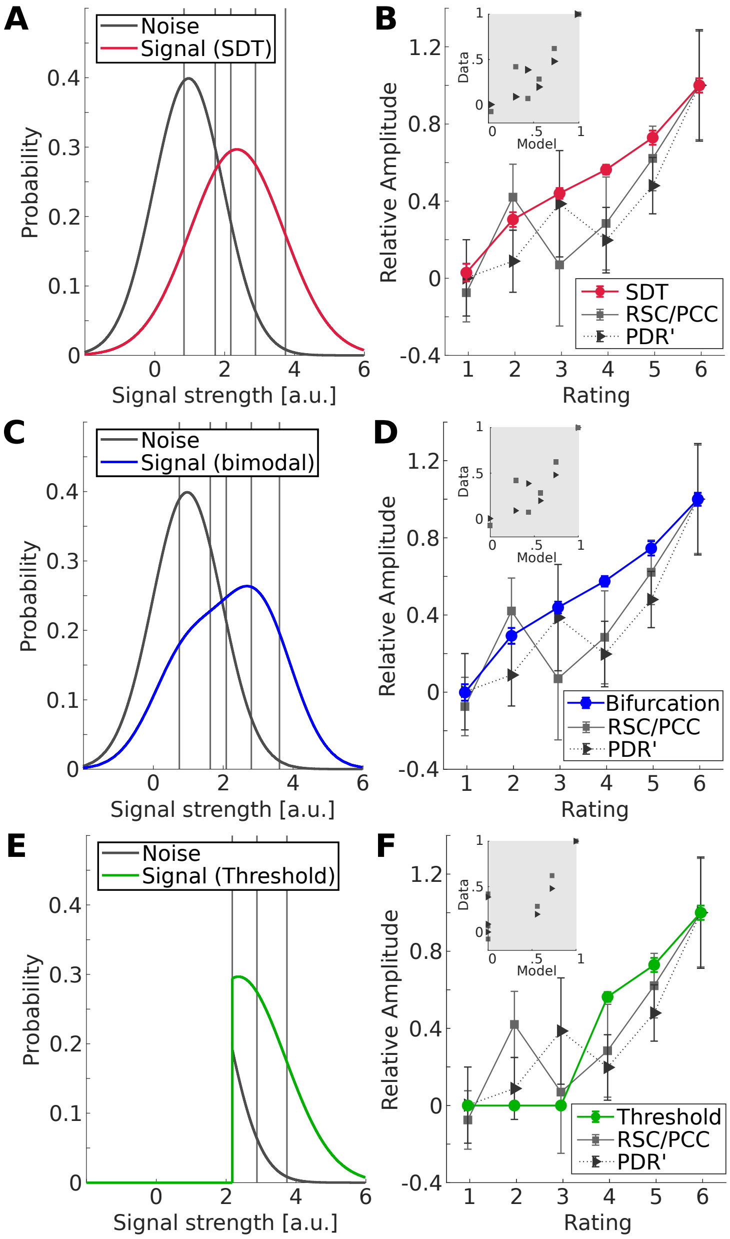


**Fig. S4 Modeling of RSC/PCC and PDR' amplitudes based on behavioral results of Experiment 2 (Rating task).** (A) Signal (red) and noise (black) distribution fitted with a STD-Model based on the behavioral results. Vertical lines represent the group average of the five criteria calculated based on the two distributions. (B) Prediction of relative amplitudes (solid line/circles) and actual amplitudes of RSC/PCC activity (solid gray line and squares) and PDR' (dashed black line and triangles). All data points are averages across participants, error bars represent the standard error of the mean. The explained variance by the model is 0.67 and 0.63 for RSC/PCC and PDR', respectively. The inset shows predicted vs. measured amplitude for RSC/PCC (gray squares) and PDR' (black triangles). (C) and (D) are the equivalent plots for a model with a bimodal signal distribution[6]. (explained variance: 0.66 for RSC/PCC, and 0.46 for PDR'), (E) and (F) for a threshold model, which assumes no AC activity below threshold (explained variance: 0.66 for RSC/PCC, 0.61 for PDR'.
